# Absence of electron transfer-associated changes in the time-dependent X-ray free-electron laser structures of the photosynthetic reaction center

**DOI:** 10.1101/2023.05.31.543167

**Authors:** Gai Nishikawa, Yu Sugo, Keisuke Saito, Hiroshi Ishikita

## Abstract

Using the X-ray free-electron laser (XFEL) structures of the photosynthetic reaction center from *Blastochloris viridis* that show light-induced time-dependent structural changes [Dods, R.et al. (2021) Nature *589*, 310-314], we investigated time-dependent changes in the energetics of the electron transfer pathway, considering the entire protein environment of the protein structures and titrating the redox active sites in the presence of all fully equilibrated titratable residues. In the dark and charge-separation intermediate structures, the calculated redox potential (*E*_m_) values for the accessory bacteriochlorophyll and bacteriopheophytin in the electron-transfer active branch (B_L_ and H_L_) are higher than those in the electron-transfer inactive branch (B_M_ and H_M_). However, the stabilization of the [P_L_P_M_]^•+^H_L_^•–^ state owing to protein reorganization is not clearly observed in the *E*_m_(H_L_) values in the charge-separated 5-ps ([P_L_P_M_]^•+^H_L_^•–^ state) structure. Furthermore, the expected chlorin ring deformation upon formation of H_L_^•–^(saddling mode) is absent in the H_L_ geometry of the original 5-ps structure. These findings suggest that there is no clear link between the time-dependent structural changes and the electron transfer events in the XFEL structures.

## INTRODUCTION

Photosynthetic reaction centers from purple bacteria (PbRC) are heterodimeric reaction centers, which are formed by the protein subunits L and M (Figure 1). In PbRC from *Blastochloris viridis*, the electronic excitation of the bacteriochlorophyll *b* (BChl*b*) pair, [P_L_P_M_], leads to electron transfer to accessory BChl*b*, B_L_, followed by electron transfer via bacteriopheophytin *b* (BPheo*b*), H_L_, to menaquinone, Q_A_, along the electron-transfer active L branch (A branch) ^1^. Electron transfer further proceeds from Q_A_ to ubiquinone, Q_B_, which is coupled with proton transfer via charged and polar residues in the Q_B_ binding region ^2^. Although the counterpart M branch (B branch) is essentially electron-transfer inactive, mutations of the Phe-L181/Tyr-M208 pair to tyrosine/phenylalanine lead to an increase in the yield of [P_L_P_M_]^•+^H_M_^•-^ formation (∼30%), which suggests that these residues are responsible for the energetic asymmetry in the electron transfer branches (e.g., ^3^). The anionic states B_L_ ^•-^, H_L_^•-^, and Q_A_^•-^ form in ∼3.5 ps, ∼5 ps, and ∼200 ps upon the formation of the electronically excited [P_L_P_M_]* state, respectively ^4^. The anionic state formation induces not only reoriganization of the protein environment ^5^ but also out-of-plane distortion of the chlorin ring ^6^. Indeed, two distinct conformations of H_L_^•-^ were reported in spectroscopic studies of PbRC from *Rhodobacter sphaeroides* ^7^.

**Figure 1.**
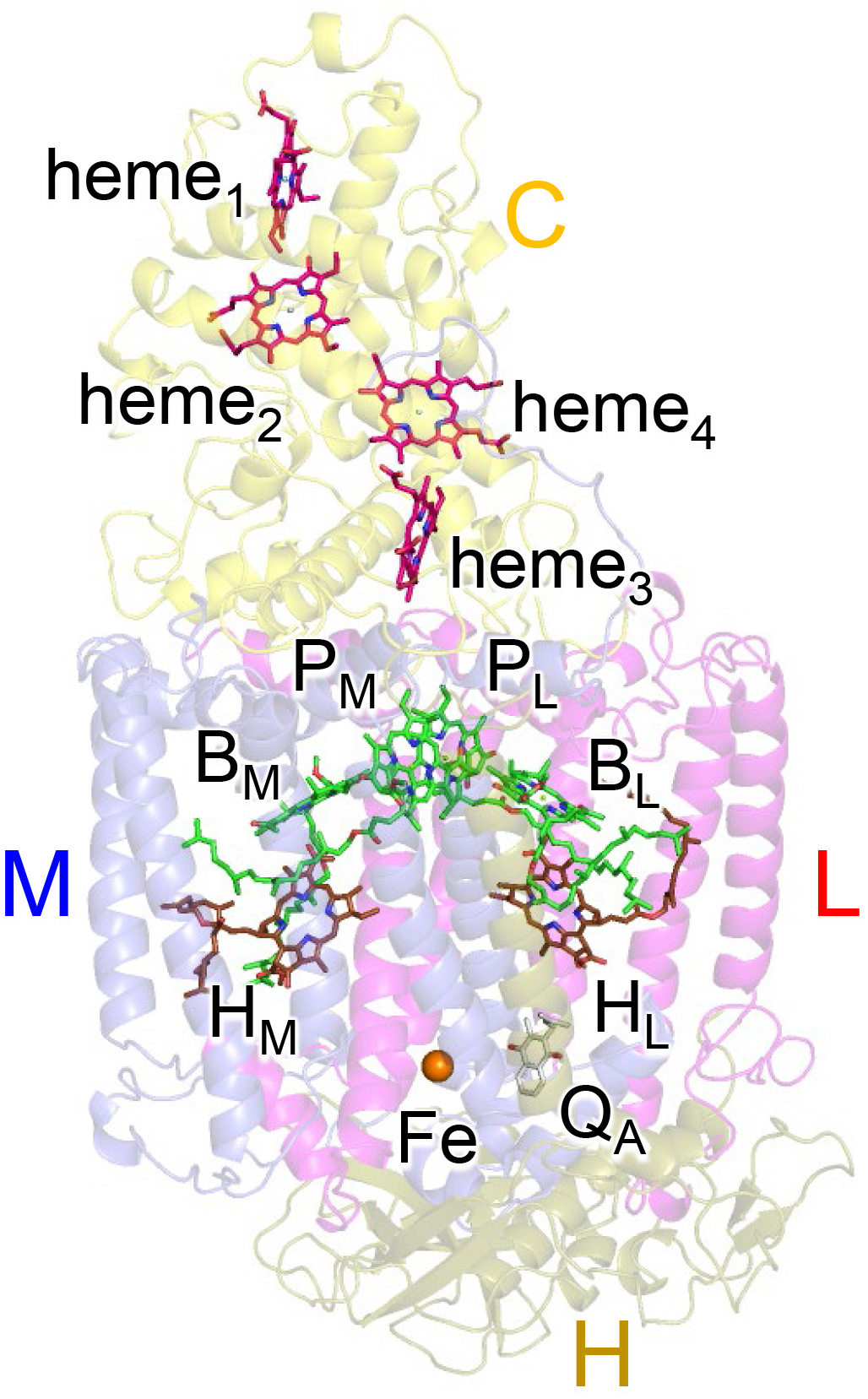
Electron transfer pathways along the L- and M-branches in PbRC from *Blastochloris viridis*. The PbRC is composed of the L (red), M (blue), H (gold), and C (yellow) subunits. [P_L_P_M_]: BChl*b* pair; B_L_ and B_M_: accessory BChl*b*; H_L_ and H_M_: BPheo*b*; Q_A_: primary quinone (menaquinone); Fe: non-heme Fe complex.

Recently, using the X-ray free electron laser (XFEL), light-induced electron density changes and structural changes of PbRC were analyzed at 1 ps, 5 ps, 20 ps, 300 ps, and 8 μs upon the electronic excitation of [P_L_P_M_] at 960 nm ^8^: the 1 ps XFEL structure represents the [P_L_P_M_]* state, the 5 ps and 20 ps XFEL structures represent the charge-separated [P_L_P_M_]^•+^H_L_^•–^ state, and the 300 ps and 8 μs XFEL structures represent the charge-separated [P_L_P_M_]^•+^Q_A_^•–^ state. According to Dods et al. ^8^, these XFEL structures revealed how the charge-separation process was stabilized by protein conformational dynamics. However, the conclusions drawn from these XFEL structures are based on data with limited resolution. Specifically, 8 out of 9 XFEL structures have a resolution of 2.8 Å (atomic coordinates from PDB codes: 5O4C, 6ZI4, and 6ZI5 for dataset a and 6ZHW, 6ZID, 6ZI6, 6ZI9, and 6ZIA for dataset b) ^8^. The data statistics may indicate that the high-resolution range of some XFEL datasets exhibits high levels of noise (e.g., low CC_1/2_). These observations raise concerns about the reliable comparison of subtle conformational changes among these XFEL structures. Hence, caution must be exercised when interpreting these XFEL structures in terms of their ability to accurately capture relevant conformational changes.

Here, we investigated how the redox potential (*E*_m_) values of the BChl*b* and BPheo*b* cofactors for one-electron reduction change as electron transfer proceeds using the dark (0 ps), 1 ps, 5 ps, 20 ps, 300 ps, and 8 μs XFEL structures, solving the linear Poisson-Boltzmann equation, and considering the protonation states of all titratable sites in the entire protein. Structural changes (e.g., side-chain orientation) in the protein environment can be analyzed in the *E*_m_ shift, as *E*_m_ is predominantly determined by the sum of the electrostatic interactions between the redox-active site and all other groups (i.e., residues and cofactors) in the protein structure. Subtle structural changes of the BChl*b* and BPheo*b* chlorin rings, which may not be pronounced even in the *E*_m_ shift ^6^, can be analyzed in the out-of-plane distortion of the chlorin rings using a normal-coordinate structural decomposition (NSD) analysis ^9, 10^ with a combination of a quantum mechanical/molecular mechanical (QM/MM) approach in the entire PbRC protein environment.

## METHODS

### Coordinates and atomic partial charges

The atomic coordinates of PbRC from *Blastochloris viridis* were taken from the XFEL structures determined at 0 ps (dark state; PDB code 5O4C for dataset a and 5NJ4 for dataset b), 1 ps ([P_L_P_M_]* state; PDB code, 6ZHW for dataset b), 5 ps ([P_L_P_M_] ^•+^H_L_^•–^ state; PDB code, 6ZI4 for dataset a and 6ZID for dataset b), 20 ps ([P_L_P_M_] ^•+^H_L_^•–^ state; PDB code, 6ZI6 for dataset b), 300 ps ([P_L_P_M_] ^•+^Q_A_^•–^ state; PDB code, 6ZI5 for dataset a and 6ZI9 for dataset b), and 8 μs ([P_L_P_M_] ^•+^Q_A_^•-^ state; PDB code, 6ZIA for dataset b). Atoms with 30% occupancy for the photoactivated state ^8^ were used wherever present. Hydrogen atoms were generated and energetically optimized with CHARMM ^11^. The atomic partial charges of the amino acids were obtained from the all-atom CHARMM22 ^12^ parameter set. For diacylglycerol, the Fe complex ^13^, and menaquinone ^14^, the atomic charges were adopted from previous studies. The atomic charges of BChl*b* and BPheo*b* (BChl*b*, BChl*b*^•+^, BChl*b*^•–^, BPheo*b*, and BPheo*b*^•–^) were determined by fitting the electrostatic potential in the neighborhood of these molecules using the RESP procedure ^15^ (Tables S1). The electronic densities were calculated after geometry optimization using the DFT method with the B3LYP functional and 6-31G** basis sets in the JAGUAR program ^16^. For the atomic charges of the nonpolar CH*_n_* groups in the cofactors (e.g., the phytol chains of BChl*b* and BPheo*b* and the isoprene side chains of quinone), a value of +0.09 was assigned to nonpolar H atoms.

### Calculation of *E*_m_: solving the linear Poisson-Boltzmann equation

The *E*_m_ values in the protein were determined by calculating the electrostatic energy difference between the two redox states in a reference model system. This was achieved by solving the linear Poisson-Boltzmann equation with the MEAD program ^17^ and using *E*_m_(BChl*b*) = –665 mV and *E*_m_(BPheo*b*) = –429 mV (based on *E*_m_(BChl*b*) = –700 mV and *E*_m_(BPheo*b*) = –500 mV for one-electron reduction measured in dimethylformamide ^18,19^), considering the solvation energy difference). The *E*_m_(Q_A_) value was calculated, using the reference *E*_m_ value of –256 mV versus NHE for menaquinone-2 in water ^20^. The difference in the *E*_m_ value of the protein relative to the reference system was added to the known *E*_m_ value. To account for the ensemble of protonation patterns, a Monte Carlo method with Karlsberg was used for sampling ^21^. The linear Poisson-Boltzmann equation was solved using a three-step grid-focusing procedure with resolutions of 2.5 Å, 1.0 Å, and 0.3 Å. Monte Carlo sampling provided the probabilities [*A_ox_*] and [*A_red_*] of the two redox states of molecule *A*, and *E*_m_ was evaluated using the Nernst equation. A bias potential was applied to ensure an equal amount of both redox states ([*A_ox_*] = [*A_red_*]), thus determining the redox midpoint potential as the resulting bias potential. To ensure consistency with previous computational results, we used identical computational conditions and parameters as previous studies (e.g., ^13^), performing all computations at 300 K, pH 7.0, and an ionic strength of 100 mM. The dielectric constants were set to 4 for the protein interior and 80 for water.

### QM/MM calculations

We employed the restricted DFT method for describing the closed-shell electronic structure and the unrestricted DFT method for the open-shell electronic structure with the B3LYP functional and LACVP* basis sets using the QSite ^22^ program. To neutralize the entire system, counter ions were added randomly around the protein using the Autoionize plugin in VMD ^23^. In the QM region, all atom positions were relaxed in the QM region, while the H-atom positions were relaxed in the MM region. The QM regions were defined as follows: for the BChl*b* pair [P_L_P_M_]: the side chains of the ligand residues (His-L173 and His-M200) and H-bod partners (His-L168, Tyr-M195, and Thr-L248); for accessory BChl*b*: B_L_/B_M_ and the side chain of the ligand residue (His-L153 for B_L_/His-M180 for B_M_); for BPheo*b*: H_L_/H_M_.

### NSD analysis

To analyze the out-of-plane distortions of chlorin rings, we employed an NSD procedure with the minimal basis approximation, where the deformation profile can be represented by the six lowest-frequency normal modes, i.e., ruffling (B_1u_), saddling (B_2u_), doming (A_2u_), waving (E_g(x)_ and E_g(y)_), and propellering (A_1u_) modes ^9, 10^. The NSD analysis was performed in the following three steps, as performed previously ^6^. First, the atomic coordinates of the Mg-substituted macrocycle were extracted from the crystal (or QM/MM optimized) structure. Second, the extracted coordinates were superimposed on the reference coordinates of the macrocycle. The superimposition is based on a least-square method, and the mathematical procedure is described in Ref. ^24^. Finally, the out-of-plane distortion in the superimposed coordinates was decomposed into the six lowest-frequency normal modes by the projection to the reference normal mode coordinates as

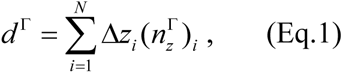

where *d*^Γ^ represents the distortion component of the mode Γ (i.e., Γ = B_1u_, B_2u_, A_2u_, E_g(x),_ E_g(y)_, or A_1u_), Δ*z_i_* is the *z*-component of the superimposed coordinates in the *i*th heavy atom, and 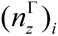 is the *z*-component of the normalized eigenvector of the reference normal mode Γ in the *i*th heavy atom. *N* represents the number of heavy atoms. See ref. ^6^ for further details.

## RESULTS AND DISCUSSION

### Energetically asymmetric electron transfer branches

The XFEL structures show that the *E*_m_ values for B_L_ are ∼50 mV higher than those for B_M_, which facilitates the formation of the charge-separated [P_L_P_M_] ^•+^B_L_^•-^ state and thereby electron transfer along the L-branch (Figures 2 and 3). As the *E*_m_ profile is substantially consistent with the *E*_m_ profile for PbRC from *Rhodobacter sphaeroides* ^13^, it seems plausible that the charge-separated [P_L_P_M_] ^•+^B_L_^•–^ and [P_L_P_M_] ^•+^H_L_^•–^ states in the active L-branch are energetically lower than the [P_L_P_M_] ^•+^B_M_^•-^ and [P_L_P_M_] ^•+^H_M_^•–^ states in the inactive M-branch, respectively, as demonstrated in QM/MM/PCM calculations ^25^. Indeed, the calculated *E*_m_ values are largely correlated with the LUMO levels calculated using a QM/MM approach, as suggested previously (coefficient of determination *R*^2^ = 0.98, Figure S1). The *E*_m_(H_L_) value of –597 mV is in line with the experimentally estimated value of ca. –600 mV for H_L_ in PbRC from *Blastochloris viridis* ^26^.

**Figure 2.**
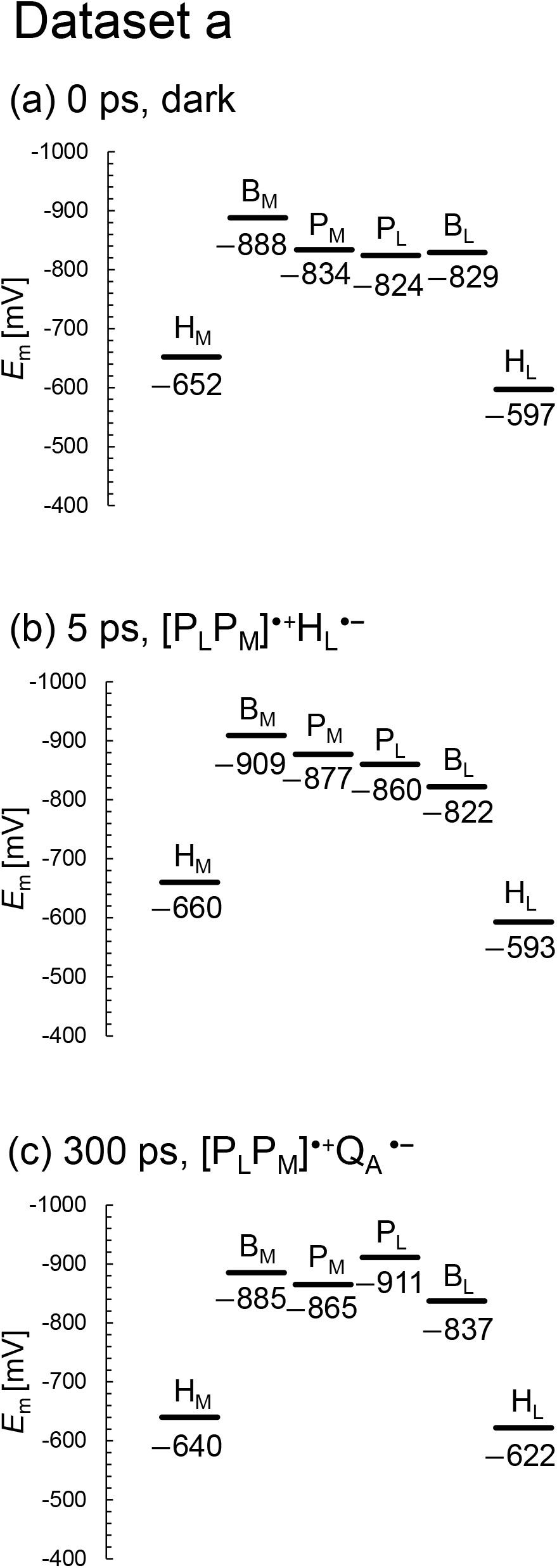
*E*_m_ profiles along the L- and M-branches in the XFEL structures for dataset a. (a) 0 ps. (b) 5 ps. (c) 300 ps.

**Figure 3.**
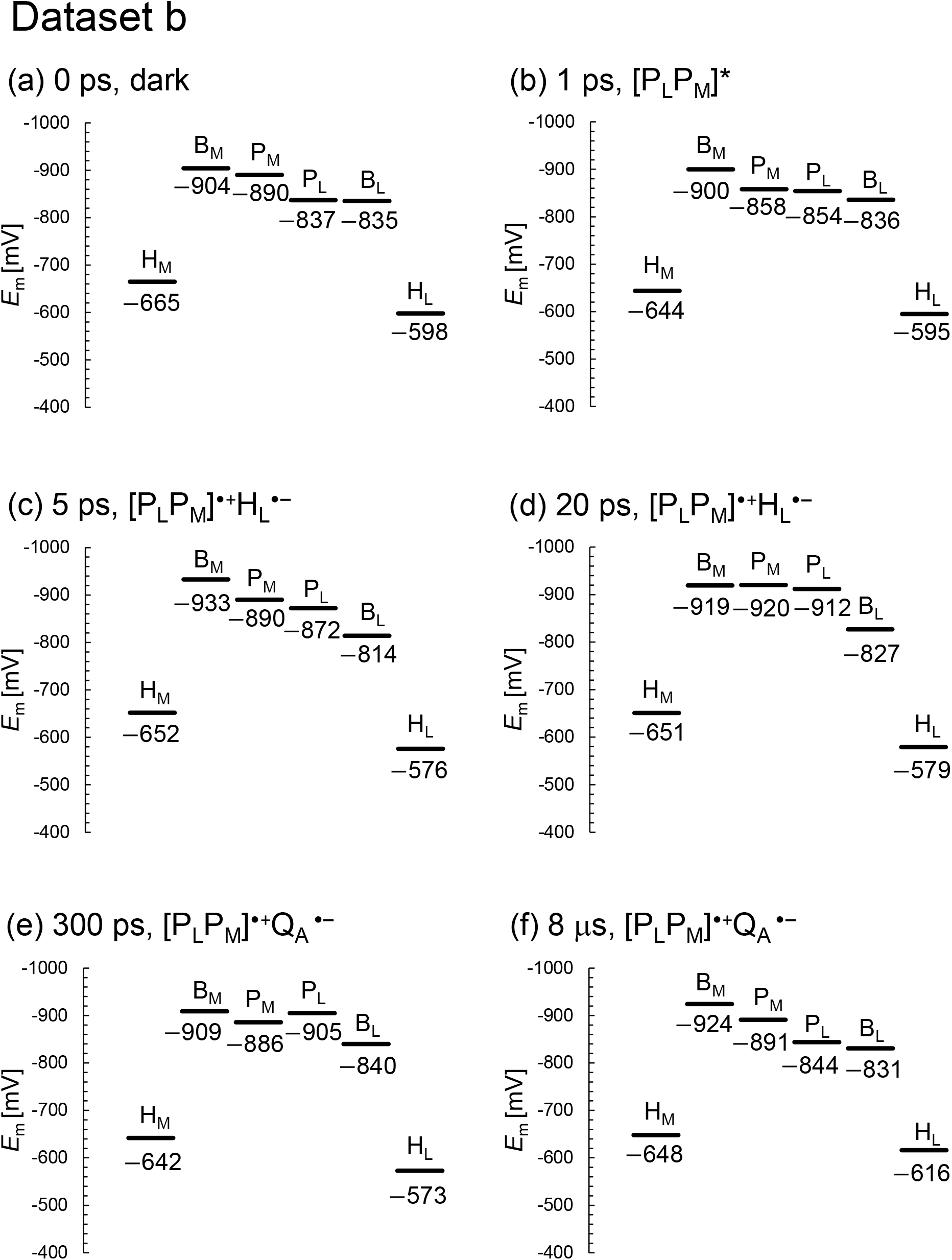
*E*_m_ profiles along the L- and M-branches in the XFEL structures for dataset b. (a) 0 ps. (b) 1 ps. (c) 5 ps. (d) 20 ps. (e) 300 ps. (f) 8 μs.

Among the L/M residue pairs, the Phe-L181/Tyr-M208 pair contributes to *E*_m_(B_L_) > *E*_m_(B_M_) most significantly (27 mV), facilitating L branch electron transfer, as suggested in theoretical studies ^27^ (Table 1, Figure 3a). This result is also consistent with the contribution of the Phe-L181/Tyr-M210 pair to the difference between *E*_m_(B_L_) and *E*_m_(B_M_), which was the largest in PbRC from *Rhodobacter sphaeroides* ^28^ (26 mV ^13^). The Asn-L158/Thr-M185 pair also contributes to the difference between *E*_m_(B_L_) and *E*_m_(B_M_) (11 mV, Table 1), as does the Val-L157/Thr-M186 pair in PbRC from *Rhodobacter sphaeroides* (22 mV ^13^).

**Table 1.** Contributions of the L/M residue pairs that are responsible for *E*_m_(B_L_) > *E*_m_(B_M_) (more than 10 mV) in the dark-state structure (mV). Difference: [contribution of subunit L to *E*_m_(B_L_)] + [contribution of subunit M to *E*_m_(B_L_)] – [contribution of subunit L to *E*_m_(B_M_)] – [contribution of subunit M to *E*_m_(B_M_)].

| Subunit L | $E_m(B_L)$ | $E_m(B_M)$ | Subunit M | $E_m(B_L)$ | $E_m(B_M)$ | Difference |
| --- | --- | --- | --- | --- | --- | --- |
| Phe-L181 | 0 | 17 | Tyr-M208 | 39 | -3 | 25 |
| His-L144 | -8 | -2 | Glu-M171 | -14 | -45 | 25 |
| Asn-L158 | 5 | -6 | Thr-M185 | -3 | -4 | 12 |

The *E*_m_ values for H_L_ are ∼50 mV higher than those for H_M_ in the dark state and [5 ps and 300 ps] XFEL structures, as observed in *E*_m_(B_L_) and *E*_m_(B_M_) (Figure 2a,c,e). However, the *E*_m_ difference decreases to ∼25 mV in the [1 ps, 20 ps, and 8 μs] XFEL structures (Figure 2b,d,f), which implies that the dark state and [5 ps and 300 ps] XFEL structures are distinct from the [1 ps, 20 ps, and 8 μs] XFEL structures (see below). Below, we discuss the dark state structure if not otherwise specified.

The Ala-L120/Asn-M147 pair contributes to *E*_m_(H_L_) > *E*_m_(H_M_) most significantly (38 mV) (Table 2, Figure S2). However, this holds true only for PbRC from *Blastochloris viridis*, as Asn-M147 is replaced with alanine (Ala-M149) in PbRC from *Rhodobacter sphaeroides*. The Asp-L218/Trp-M252 pair decreases *E*_m_(H_M_) with respect to *E*_m_(H_L_), thereby contributing to *E*_m_(H_L_) > *E*_m_(H_M_) (20 mV) (Table 2, Figure S2). Arg-L103 orients toward the protein interior, whereas Arg-M130 orients toward the protein exterior (Figure S2), which contributes to *E*_m_(H_L_) > *E*_m_(H_M_) (17 mV) (Table 2). Ser-M271 forms an H-bond with Asn-M147 near H_M_ (Figure 3b). Thus, the contribution of Ser-M271 to *E*_m_(H_L_) is large, although this residue is replaced with alanine (Ala-M273) in PbRC from *Rhodobacter sphaeroides*.

**Table 2.** Contributions of the L/M residue pairs that are responsible for *E*_m_(H_L_) > *E*_m_(H_M_) (more than 10 mV) in the dark-state structure (mV). Difference: [contribution of subunit L to *E*_m_(H_L_)] + [contribution of subunit M to *E*_m_(H_L_)] – [contribution of subunit L to *E*_m_(H_M_)] – [contribution of subunit M to *E*_m_(H_M_)].

| Subunit L | $E_m(H_L)$ | $E_m(H_M)$ | Subunit M | $E_m(H_L)$ | $E_m(H_M)$ | Difference |
| --- | --- | --- | --- | --- | --- | --- |
| Ala-L120 | -4 | 0 | Asn-M147 | 0 | -42 | 38 |
| Asp-L218 | -2 | -22 | Trp-M252 | 1 | 0 | 20 |
| Arg-L103 | 77 | 3 | Arg-M130 | 3 | 59 | 17 |
| Ala-L237 | -2 | 0 | Ser-M271 | 3 | -16 | 16 |
| Lys-L110 | 17 | 2 | Ala-M137 | 0 | 3 | 14 |
| Val-L219 | 1 | 5 | Thr-M253 | 17 | 1 | 11 |
| His-L211 | 1 | 0 | Arg-M245 | 14 | 4 | 11 |

### Relevance of structural changes observed in XFEL structures

According to Dods et al., the 5-ps and 20-ps structures correspond to the charge-separated [P_L_P_M_]^•+^H_L_^•–^ state ^8^. If this is the case, *E*_m_(H_L_) is expected to be exclusively higher in the 5-ps and 20-ps structures than in the other XFEL structures due to the stabilization of the [P_L_P_M_]^•+^H_L_^•–^ state by protein reorganization. In dataset a, the *E*_m_(H_L_) value is only 4 mV higher in the 5-ps structure than in the dark structure (Figure 5a). In dataset b, the *E*_m_(H_L_) value is ∼20 mV higher in the 5- and 20-ps structures than in the dark structure (Figure 5b). However, the *E*_m_(H_L_) value is 25 mV higher in the 300-ps structure than in the dark structure. Tables 3 and 4 show the residues that contribute to the slight increase in *E*_m_(H_L_) most significantly in the 5- and 20-ps structures. Most of these residues were in the region where Dods et al. specifically performed multiple rounds of partial occupancy refinement (e.g., 153–178, 190, 230 and 236–248 of subunit L and 193–221, 232, 243– 253, 257–266 of subunit M) ^8^. In dataset b (Table 4), which has more data points than dataset a (Table 3), the contributions of these residues to *E*_m_(H_L_) often fluctuate (e.g., upshift/downshift followed by downshift/upshift) at different time intervals (e.g., 1 to 5 ps, 5 to 20 ps, and 20 to 300 ps). This result suggests that the structural differences among the XFEL structures are not related to the actual time course of charge separation. Furthermore, the *E*_m_(H_M_) value in the inactive M branch is also ∼15 mV higher in the 5- and 20-ps structures than in the dark structure (Figure 5b). These results suggest that the ∼20 mV higher *E*_m_(H_L_) value in the 5- and 20-ps structures is not specifically due to the formation of the [P_L_P_M_]^•+^H_L_^•–^ state. Thus, the stabilization of the [P_L_P_M_]^•+^H_L_^•–^ state owing to protein reorganization is not clearly observed in the *E*_m_(H_L_) values.

**Figure 4.**
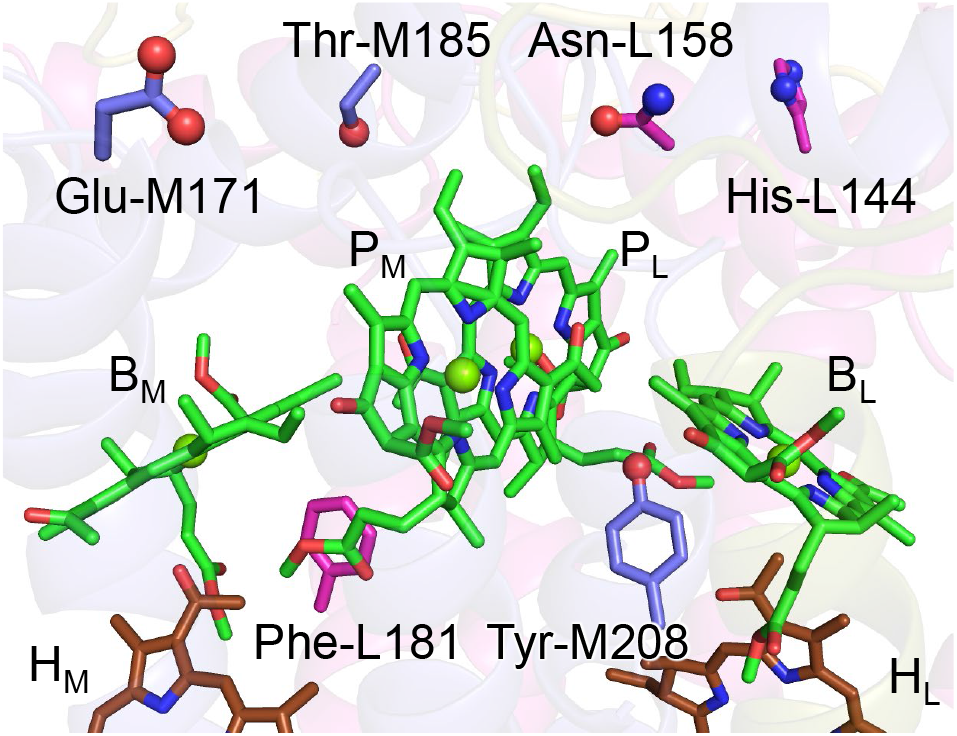
Residue pairs that are responsible for *E*_m_(B_L_) > *E*_m_(B_M_).

**Figure 5.**
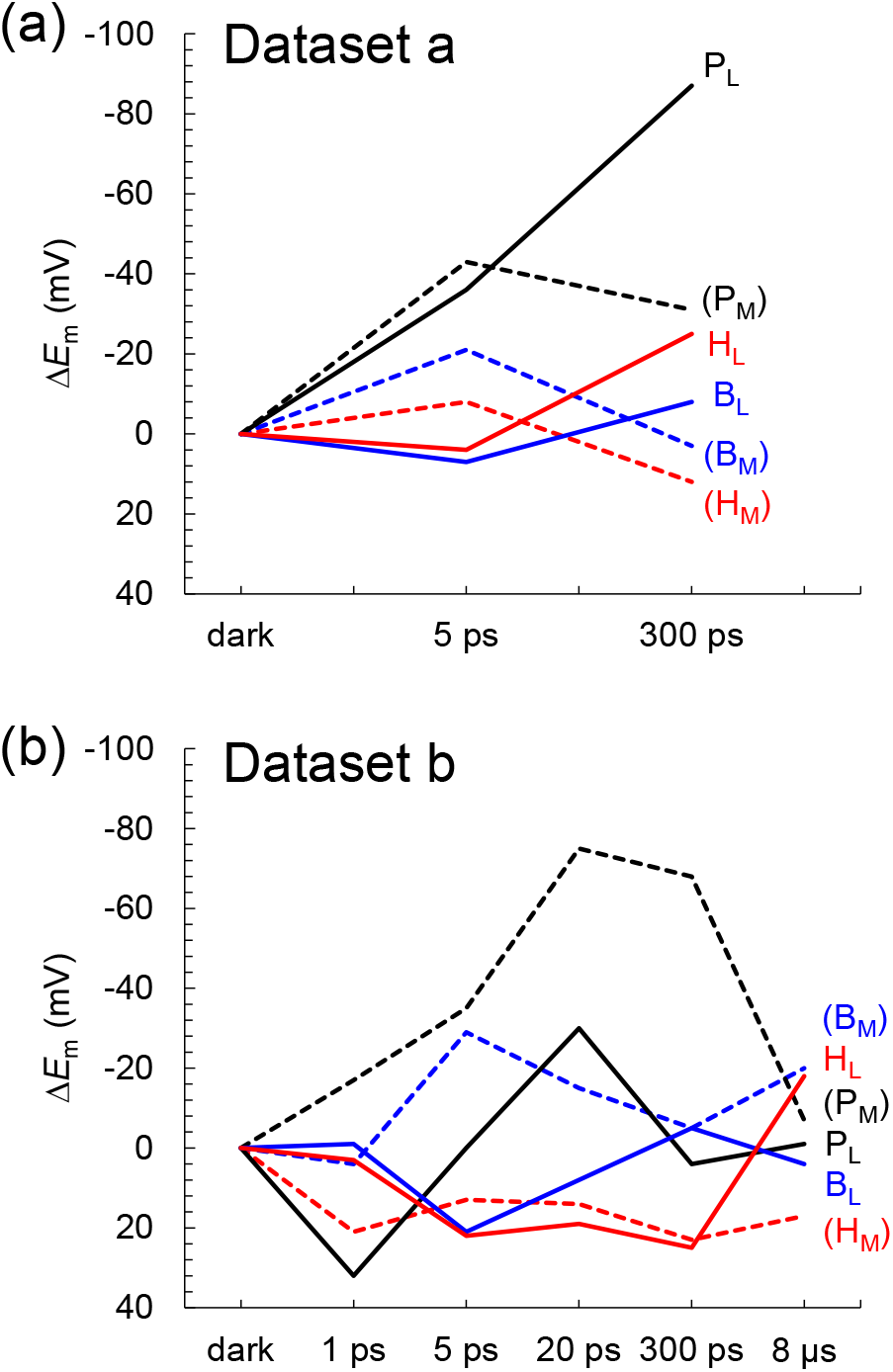
Time-dependent *E*_m_ changes for BChl*b* and BPheo*b* in the XFEL structures. (a) Dataset a. (b) Dataset b. Δ*E*_m_ denotes the *E*_m_ shift with respect to the dark state structure. Black solid lines: P_L_; black dotted lines: P_M_; blue solid lines B_L_; blue dotted lines: B_M_; red solid lines: H_L_; red dotted lines: H_M_.

**Table 3.** Residues that shift *E*_m_(H_L_) most significantly during putative electron transfer in the XFEL structures (dataset a) (mV). The same residues are highlighted in the same colors for clarity.

| Dataset a |  | shift |  | shift |
| --- | --- | --- | --- | --- |
| 0 to 5 ps | Ser-L176 | 5 | Cys-M210 | 4 |
|  | Thr-M220 | -7 | B<sub>L</sub> | -5 |
| 5 to 300 ps | B<sub>L</sub> | 7 | Gly-M209 | 3 |
|  | Gly-M211 | -11 | Leu-M212 | -8 |

**Table 4.** Residues that shift *E*_m_(H_L_) most significantly during putative electron transfer in the XFEL structures (dataset b) (mV). The same residues are highlighted in the same colors for clarity.

| Dataset b |  | shift |  | shift |
| --- | --- | --- | --- | --- |
| 0 to 1 ps | Ser-L238 | 8 | Ser-L176 | 7 |
|  | B<sub>L</sub> | -7 | Leu-M213 | -3 |
| 1 to 5 ps | Gly-M211 | 6 | Leu-M213 | 5 |
|  | Ser-L238 | -6 | Thr-M253 | -5 |
| 5 to 20 ps | B<sub>L</sub> | 12 | Thr-M253 | 7 |
|  | Leu-M213 | -4 | P<sub>M</sub> | -3 |
| 20 to 300 ps | Ser-L238 | 3 | Gly-M211 | 2 |
|  | B<sub>L</sub> | -10 | Glu-L212 | -4 |
| 300 ps to 8 μs | Glu-L212 | 4 | Leu-M213 | 4 |
|  | B<sub>L</sub> | -6 | Gly-M211 | -5 |

A normal-coordinate structural decomposition (NSD) analysis ^9, 10^ of the out-of-plane distortion of the chlorin ring is sensitive to subtle structural changes in the chlorin ring, which are not distinct in the *E*_m_ changes ^6^. QM/MM calculations indicate that H_L_^•-^ formation induces the saddling mode in the chlorin ring, which describes the movement of rings I and III being in the opposite direction to the movement of rings II and IV along the normal axis of the chlorin ring (Tables 5 and 6). However, (i) in the XFEL structures, the saddling mode of H_L_ remains practically unchanged in dataset a during electron transfer (Figure 6 and Tables S2 and S3). In dataset b, the saddling mode of H_L_ is induced most significantly at 1 ps, which does not correspond to the charge-separated [P_L_P_M_]^•+^H_L_^•–^ state (Figure 7). (ii) In addition, the ruffling mode is more pronounced than the saddling mode in H_L_ (Figure 7), which suggests that the observed deformation of H_L_ is not directly associated with the reduction of H_L_.

**Figure 6.**
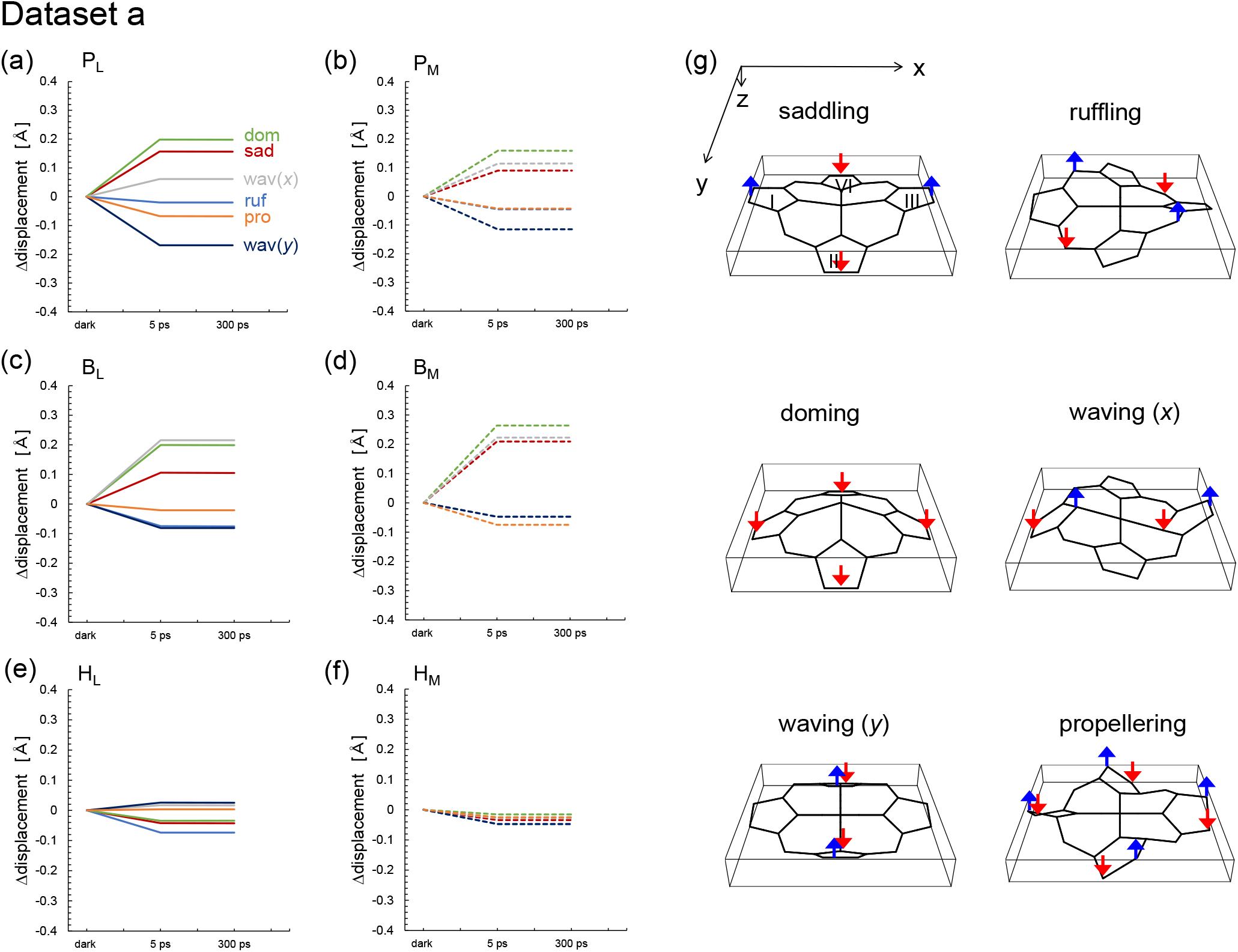
Time-dependent changes in the lowest frequency out-of-plane modes of the chlorin rings in the XFEL structures (dataset a). Sad: saddling (red); ruf: ruffling (blue); dom: doming (green); wav(*x*, *y*): waving (*x*, *y*) (gray, dark blue); pro: propellering (orange). Solid and dotted lines indicate L and M branches, respectively. See Table S2 for the absolute values in the dark state for dataset a.

**Figure 7.**
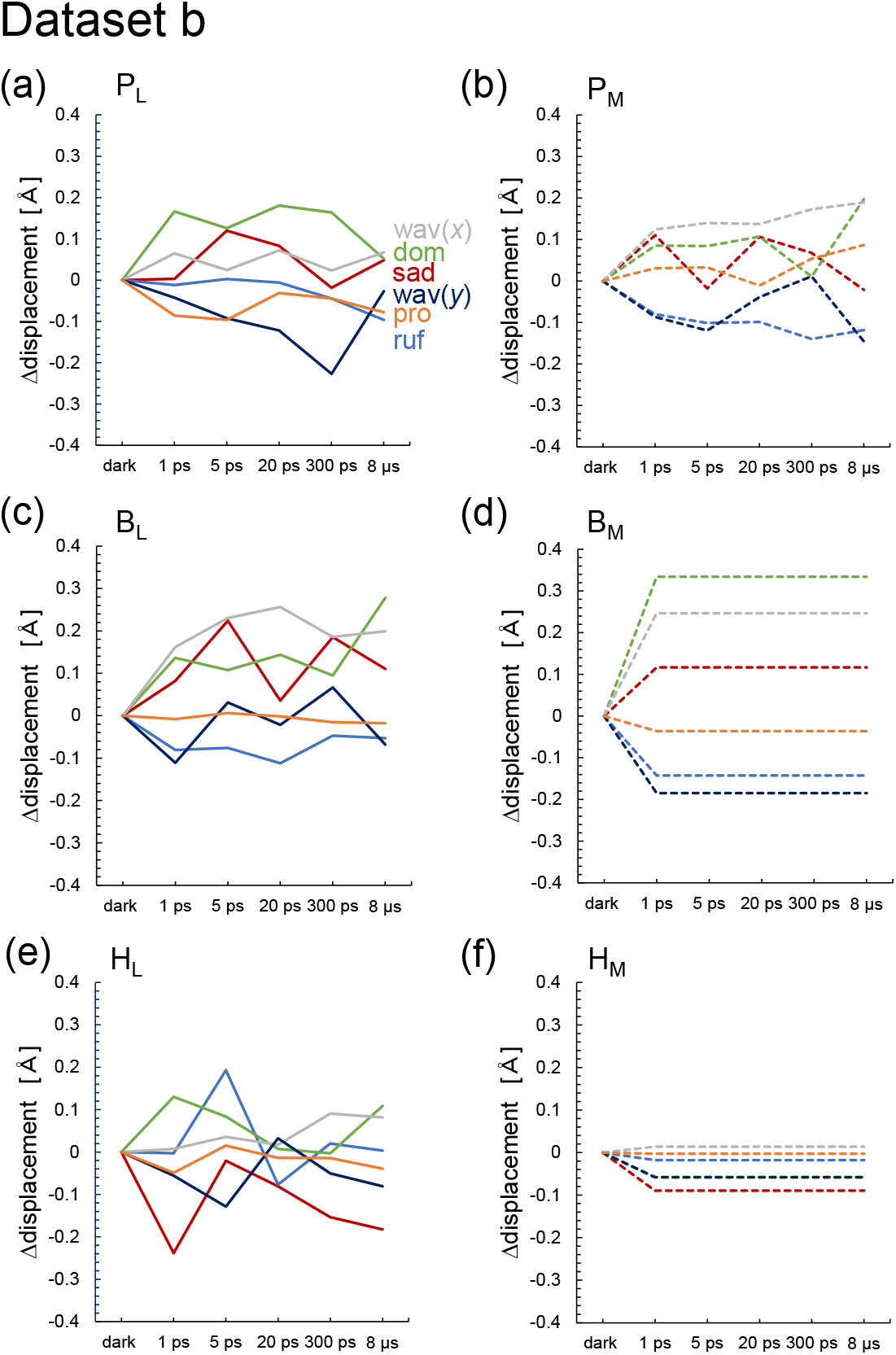
Time-dependent changes in the lowest frequency out-of-plane modes of the chlorin rings in the XFEL structures (dataset b). Sad: saddling (red); ruf: ruffling (blue); dom: doming (green); wav(*x*, *y*): waving (*x*, *y*) (gray, dark blue); pro: propellering (orange). Solid and dotted lines indicate L and M branches, respectively. See Table S3 for the absolute values in the dark state for dataset b.

**Table 5.**
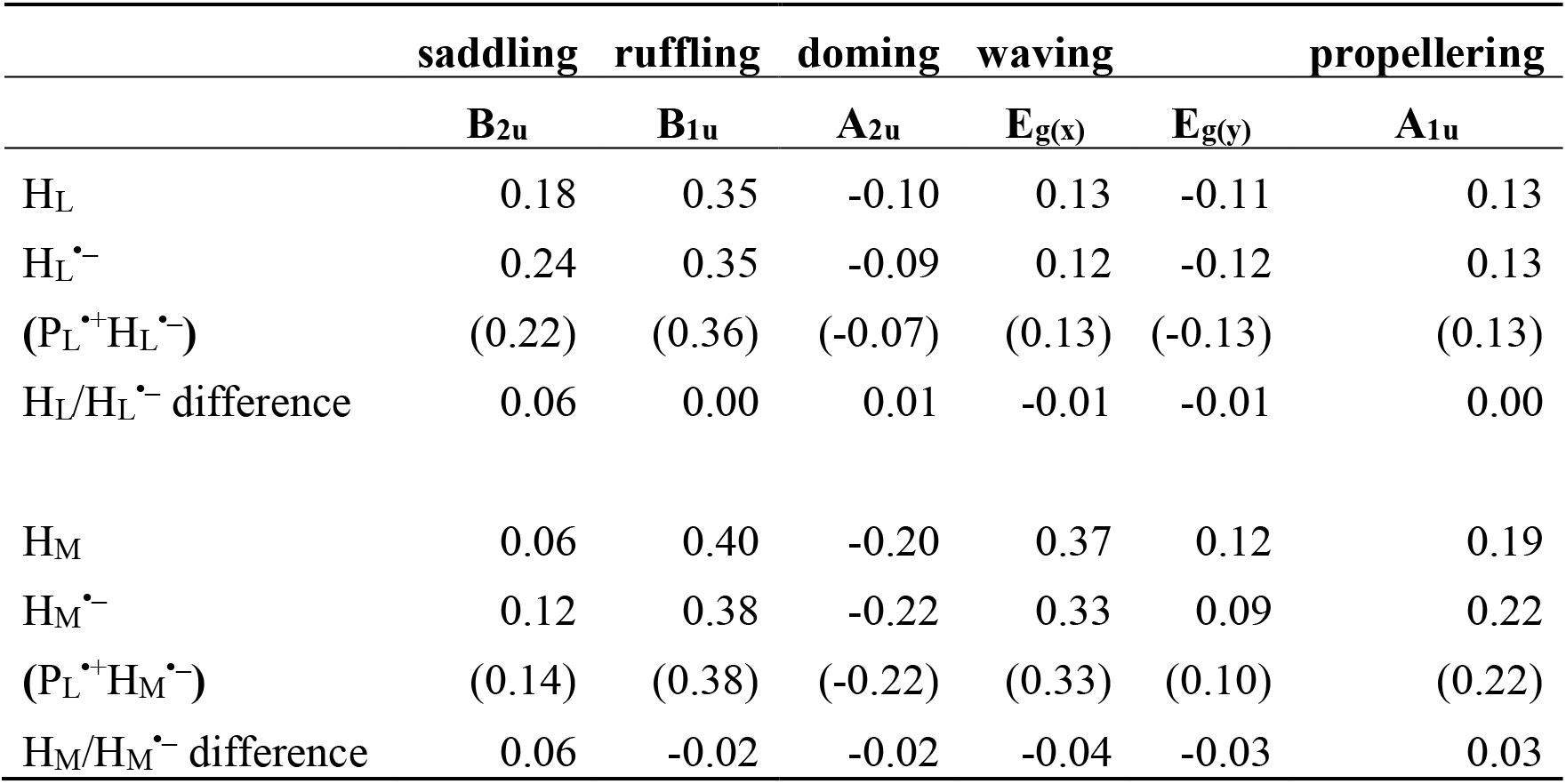
Induced out-of-plane distortion of H_L_ and H_M_ in the PbRC protein environment of the dark structure for dataset a in response to the reduction (Å).

**Table 6.**
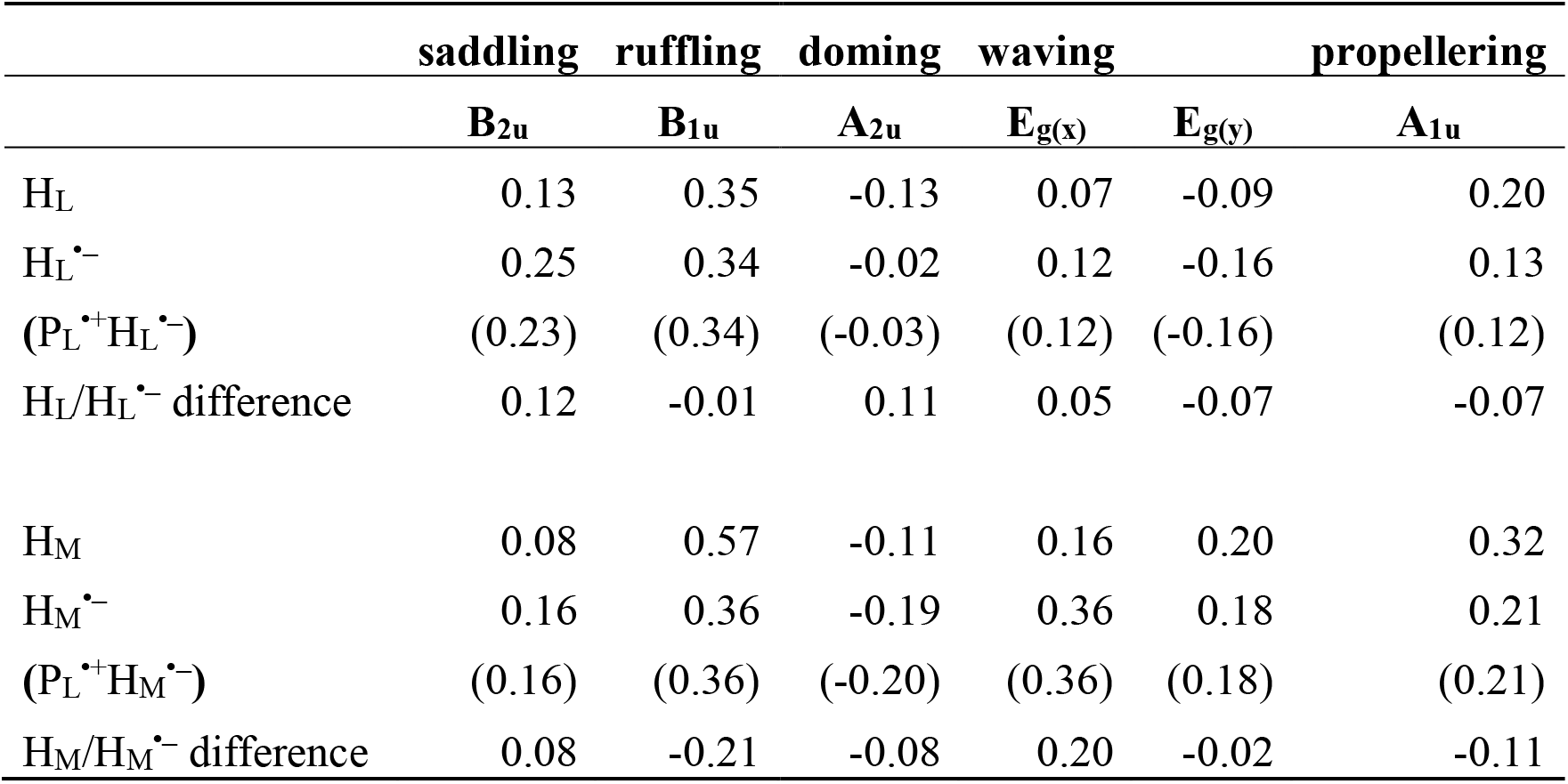
Induced out-of-plane distortion of H_L_ and H_M_ in the PbRC protein environment of the dark structure for dataset b in response to the reduction (Å).

One might argue that the loss of the link between the formation of the charge-separated state and the *E*_m_(H_L_) change (Figure 5) is not due to experimental errors but rather represents the actual ps-timescale phenomena during the primary charge-separation reactions (e.g., Dods et al. noted that “the primary electron-transfer step to H_L_ is more rapid than conventional Marcus theory” ^8^). However, even if this were the case, this hypothesis regarding the relevance of the XFEL structures to the electron-transfer events could be further explored by examining the changes in *E*_m_(Q_A_) among the XFEL structures, considering the relatively slow electron-transfer step to Q_A_ that allows sufficient protein relaxation to occur (e.g., Dods et al. stated that “the electron-transfer step to Q_A_ has a single exponential decay time of 230 ± 30 ps, consistent with conventional Marcus theory” ^8^). That is, if the *E*_m_(Q_A_) values are not higher in the 300-ps and 8-μs structures than in the other structures, it suggests that significant experimental errors exist, rendering the XFEL structures irrelevant to the electron transfer events. Consistent with this perspective, the present results demonstrate that the *E*_m_(Q_A_) values in the 300-ps and 8-μs structures are not significantly higher than those in the other structures, including the dark state structure (Figure 8). Consequently, the lack of a clear relationship between the charge separated state and the changes in *E*_m_(Q_A_) at 300 ps and 8-μs further strengthens the argument that the XFEL structures are irrelevant to the electron transfer events.

**Figure 8.**
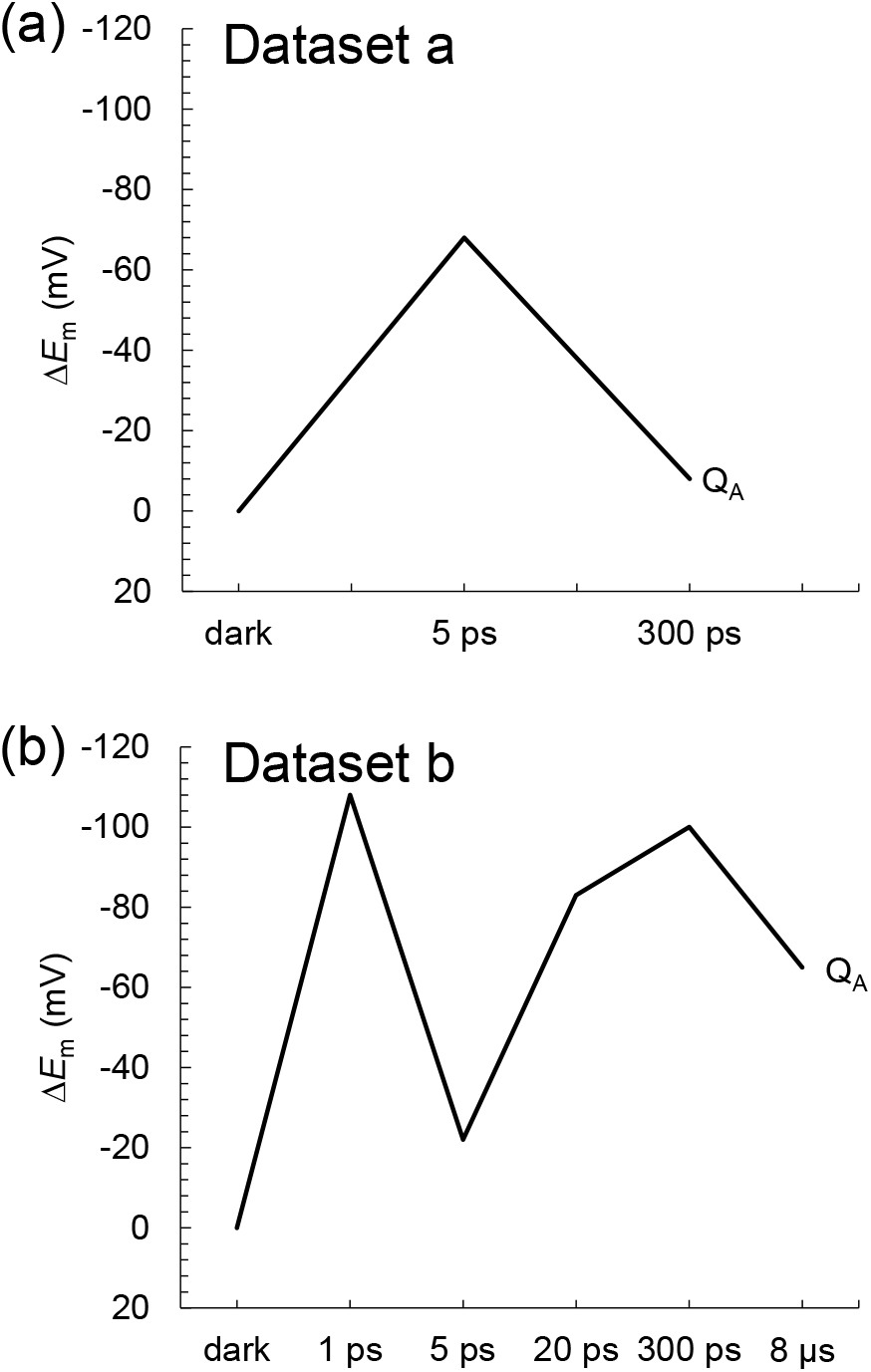
Time-dependent *E*_m_ changes for Q_A_ in the XFEL structures. (a) Dataset a. (b) Dataset b. Δ*E*_m_ denotes the *E*_m_ shift with respect to the dark state structure. Note that the calculated *E*_m_(Q_A_) values for dataset a and dataset b in the dark structure are –223 mV and –209 mV, respectively, which are comparable to experimentally measured values of –150 mV for PbRC from *Blastochloris viridis* (menaquinone) ^29^ and –180 mV for PbRC from *Rhodobacter sphaeroides* (ubiquinone) ^30^.

In summary, the *E*_m_ values in the active L branch are higher than those in the inactive M branch in the XFEL structures, which suggests that electron transfer via B_L_^•-^ and H_L_^•-^ is energetically more favored than that via B_M_^•-^ and H_M_^•–^ (Figure 2). The Phe-L181/Tyr-M208 pair contributes to the difference between *E*_m_(B_L_) and *E*_m_(B_M_) the most significantly, as observed in the Phe-L181/Tyr-M210 pair in PbRC from *Rhodobacter sphaeroides* ^13, 28^. The stabilization of the [P_L_P_M_]^•+^H_L_^•–^ state owing to protein reorganization is not clearly observed in the *E*_m_(H_L_) values (Figure 5). The absence of the induced saddling mode in the H_L_ chlorin ring in the 5- and 20-ps structures suggests that H_L_^•–^ does not specifically exist in these XFEL structures (Figures 6 and 7). The cyclic fluctuations in the contributions of the residues to *E*_m_(H_L_) at different time intervals suggest that the structural differences among the XFEL structures are not related to the actual time course of charge separation (Table 4). The major limitation of the structural studies conducted by Dods et al. ^8^ is the relatively low resolution of their XFEL structures, primarily at 2.8 Å. Consequently, the observed changes in *E*_m_ values and chlorin ring deformations are more likely to reflect experimental errors rather than actual structural changes induced by electron transfer events. This concern is reinforced by the lack of a clear relationship between the actual Q_A_^•-^ formation and the *E_m_*(Q_m_) values in the 300-ps and 8-μs structures (Figure 8). Consequently, the proposed time-dependent structural changes proposed by Dods et al. ^8^ are highly likely irrelevant to the electron transfer events.

Hence, it is crucial to exercise caution when interpreting time-dependent XFEL structures, especially in the absence of comprehensive evaluations of the energetics and accompanying structural changes. This cautionary note should serve as a counterargument in the future, highlighting the potential pitfalls associated with presenting time-dependent XFEL structures of insufficient quality and drawing conclusive interpretations of protein structural changes that may not be distinguishable from significant experimental errors. Future high-resolution structures may provide further insights into the actual structural changes relevant to electron transfer events. By combining both high-resolution structures and rigorous energetic evaluations, a more comprehensive understanding of the protein structure-function relationship can be achieved.

## ACKNOWLEDGMENTS

This research was supported by JSPS KAKENHI (JP23H04963 to K.S.; JP20H03217 and JP23H02444 to H.I.) and Interdisciplinary Computational Science Program in CCS, University of Tsukuba.

