## Supplementary material for "Absence of electron transfer-associated changes in the time-dependent X-ray free-electron laser structures of the photosynthetic reaction center": Figures S1 and S2, and Tables S1 to S3

**Contents**

8 pages

2 Figures

3 tables

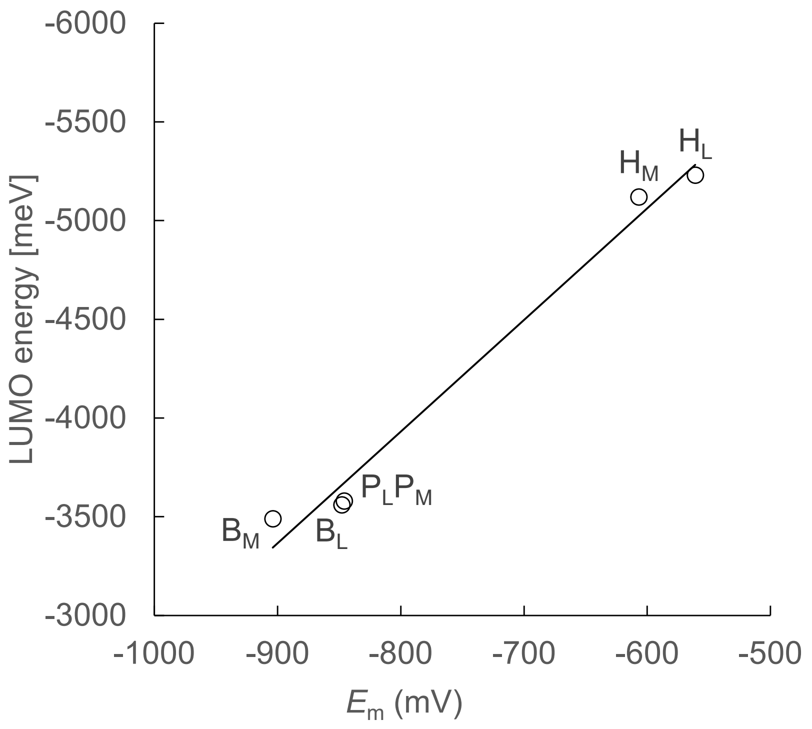


**Figure S1.** $E_m$ values calculated solving the linear Poisson-Boltzmann equation and LUMO energy levels calculated using a QM/MM approach in the dark-state structure.

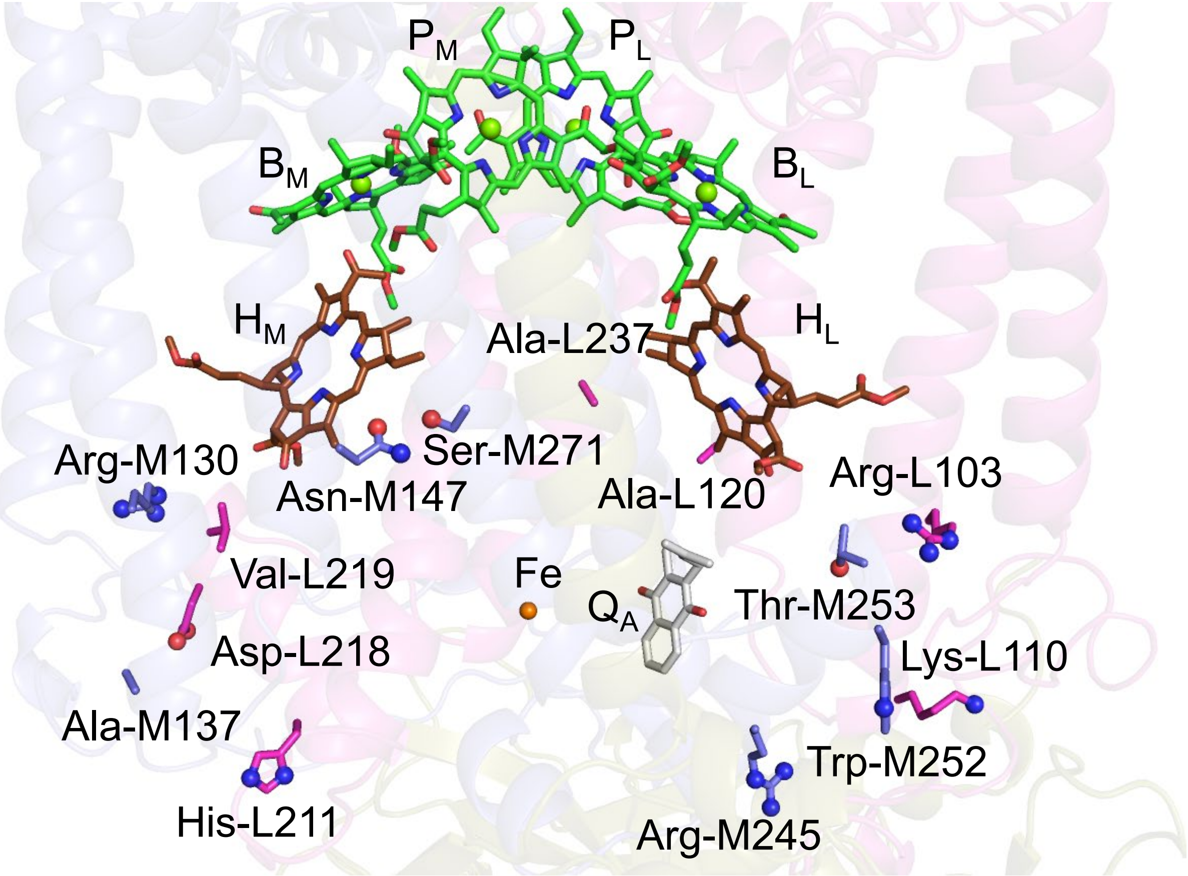


**Figure S2.** Residue pairs that are responsible for $E_m(H_L) > E_m(H_M)$.

**Table S1.** Atomic charges of BChl*b* and BPheo*b*.

| | **BChl*b*$^{•+}$** | **BChl*b*** | **BChl*b*$^{•-}$** | **BPheo*b*** | **BPheo*b*$^{•-}$** |
|---|---|---|---|---|---|
| MG | 0.889 | 0.851 | 0.795 | | |
| CHA | -0.038 | 0.071 | 0.081 | 0.118 | 0.090 |
| CHB | -0.453 | -0.400 | -0.410 | -0.354 | -0.392 |
| HHB | 0.168 | 0.154 | 0.138 | 0.156 | 0.150 |
| CHC | -0.427 | -0.389 | -0.432 | -0.314 | -0.352 |
| HHC | 0.189 | 0.171 | 0.163 | 0.107 | 0.088 |
| CHD | -0.414 | -0.402 | -0.405 | -0.331 | -0.369 |
| HHD | 0.175 | 0.169 | 0.158 | 0.160 | 0.155 |
| NA | -0.353 | -0.358 | -0.358 | -0.236 | -0.242 |
| C1A | 0.124 | 0.041 | -0.024 | -0.083 | -0.136 |
| C2A | -0.012 | 0.045 | 0.067 | 0.231 | 0.260 |
| H2A | 0.075 | 0.056 | 0.050 | 0.020 | 0.012 |
| C3A | 0.116 | 0.065 | 0.052 | 0.153 | 0.161 |
| H3A | 0.049 | 0.055 | 0.042 | 0.012 | -0.011 |
| C4A | 0.301 | 0.263 | 0.235 | 0.159 | 0.117 |
| CMA | -0.283 | -0.456 | -0.494 | -0.429 | -0.397 |
| HMA1 | 0.086 | 0.124 | 0.126 | 0.108 | 0.090 |
| HMA2 | 0.086 | 0.124 | 0.126 | 0.108 | 0.090 |
| HMA3 | 0.086 | 0.124 | 0.126 | 0.108 | 0.090 |
| CAA | -0.062 | 0.049 | 0.033 | -0.028 | 0.023 |
| HAA1 | 0.073 | 0.000 | 0.000 | 0.036 | 0.020 |
| HAA2 | 0.073 | 0.000 | 0.000 | 0.036 | 0.020 |
| CBA | -0.498 | -0.296 | -0.308 | -0.549 | -0.575 |
| HBA1 | 0.140 | 0.069 | 0.082 | 0.145 | 0.135 |
| HBA2 | 0.140 | 0.069 | 0.082 | 0.145 | 0.135 |
| CGA | 0.821 | 0.771 | 0.782 | 0.853 | 0.873 |
| O1A | -0.527 | -0.517 | -0.533 | -0.529 | -0.535 |
| O2A | -0.391 | -0.400 | -0.409 | -0.411 | -0.417 |
| NB | -0.517 | -0.458 | -0.424 | -0.213 | -0.172 |
| HNB | | | | 0.222 | 0.204 |
| C1B | 0.301 | 0.180 | 0.117 | 0.132 | 0.092 |
| C2B | 0.162 | 0.178 | 0.168 | 0.190 | 0.150 |
| C3B | -0.372 | -0.435 | -0.525 | -0.378 | -0.467 |
| C4B | 0.432 | 0.368 | 0.387 | 0.232 | 0.237 |
| CMB | -0.375 | -0.326 | -0.289 | -0.392 | -0.277 |
| HMB1 | 0.130 | 0.099 | 0.073 | 0.125 | 0.076 |
| HMB2 | 0.130 | 0.099 | 0.073 | 0.125 | 0.076 |

| | | | | | | |
|---|---|---|---|---|---|---|
| HMB3 | 0.130 | 0.099 | 0.073 | | 0.125 | 0.076 |
| CAB | 0.684 | 0.714 | 0.732 | | 0.685 | 0.713 |
| OBB | -0.466 | -0.503 | -0.557 | | -0.488 | -0.545 |
| CBB | -0.521 | -0.541 | -0.540 | | -0.399 | -0.428 |
| HBB1 | 0.154 | 0.144 | 0.126 | | 0.105 | 0.097 |
| HBB2 | 0.154 | 0.144 | 0.126 | | 0.105 | 0.097 |
| HBB3 | 0.154 | 0.144 | 0.126 | | 0.105 | 0.097 |
| NC | -0.332 | -0.293 | -0.300 | | -0.215 | -0.232 |
| C1C | 0.146 | 0.046 | 0.000 | | 0.031 | -0.025 |
| C2C | 0.403 | 0.448 | 0.485 | | 0.466 | 0.565 |
| H2C | 0.001 | 0.000 | 0.000 | | -0.026 | -0.062 |
| C3C | -0.320 | -0.311 | -0.295 | | -0.327 | -0.336 |
| C4C | 0.356 | 0.276 | 0.229 | | 0.260 | 0.241 |
| CMC | -0.376 | -0.369 | -0.356 | | -0.394 | -0.357 |
| HMC1 | 0.111 | 0.093 | 0.076 | | 0.098 | 0.076 |
| HMC2 | 0.111 | 0.093 | 0.076 | | 0.098 | 0.076 |
| HMC3 | 0.111 | 0.093 | 0.076 | | 0.098 | 0.076 |
| CAC | 0.042 | -0.029 | -0.075 | | -0.024 | -0.061 |
| HAC | 0.104 | 0.095 | 0.085 | | 0.092 | 0.081 |
| CBC | -0.248 | -0.216 | -0.180 | | -0.190 | -0.160 |
| HBC1 | 0.103 | 0.076 | 0.053 | | 0.070 | 0.047 |
| HBC2 | 0.103 | 0.076 | 0.053 | | 0.070 | 0.047 |
| HBC3 | 0.103 | 0.076 | 0.053 | | 0.070 | 0.047 |
| ND | -0.528 | -0.447 | -0.382 | | -0.060 | -0.030 |
| HND | | | | | 0.081 | 0.070 |
| C1D | 0.313 | 0.226 | 0.156 | | 0.075 | 0.041 |
| C2D | 0.113 | 0.128 | 0.094 | | 0.153 | 0.111 |
| C3D | -0.202 | -0.260 | -0.285 | | -0.288 | -0.325 |
| C4D | 0.253 | 0.123 | 0.079 | | 0.047 | 0.044 |
| CMD | -0.341 | -0.284 | -0.217 | | -0.290 | -0.226 |
| HMD1 | 0.128 | 0.094 | 0.059 | | 0.098 | 0.065 |
| HMD2 | 0.128 | 0.094 | 0.059 | | 0.098 | 0.065 |
| HMD3 | 0.128 | 0.094 | 0.059 | | 0.098 | 0.065 |
| CAD | 0.589 | 0.643 | 0.637 | | 0.674 | 0.661 |
| OBD | -0.420 | -0.475 | -0.537 | | -0.478 | -0.538 |
| CBD | -0.594 | -0.639 | -0.675 | | -0.641 | -0.649 |
| HBD | 0.266 | 0.230 | 0.221 | | 0.218 | 0.207 |
| CGD | 0.787 | 0.785 | 0.819 | | 0.770 | 0.780 |
| O1D | -0.497 | -0.516 | -0.547 | | -0.508 | -0.532 |
| O2D | -0.334 | -0.345 | -0.354 | | -0.338 | -0.340 |

| | | | | | | |
|---|---|---|---|---|---|---|
| CED | -0.066 | -0.001 | 0.054 | | 0.003 | 0.057 |
| HED1 | 0.098 | 0.073 | 0.047 | | 0.073 | 0.047 |
| HED2 | 0.098 | 0.073 | 0.047 | | 0.073 | 0.047 |
| HED3 | 0.098 | 0.073 | 0.047 | | 0.073 | 0.047 |
| C1 | 0.108 | 0.078 | 0.098 | | 0.100 | 0.105 |
| H1A | 0.072 | 0.070 | 0.055 | | 0.060 | 0.048 |
| H1B | 0.072 | 0.070 | 0.055 | | 0.060 | 0.048 |
| **total** | 1.000 | 0.000 | -1.000 | | 0.000 | -1.000 |

**Table S2**. Out-of-plane distortions in the PbRC protein environment of the dark structure for dataset a (Å).

| | **saddling** | **ruffling** | **doming** | **waving** | | **propellering** |
|---|---|---|---|---|---|---|
| | $\mathbf{B_{2u}}$ | $\mathbf{B_{1u}}$ | $\mathbf{A_{2u}}$ | $\mathbf{E_{g(x)}}$ | $\mathbf{E_{g(y)}}$ | $\mathbf{A_{1u}}$ |
| $P_L$ | 0.10 | -0.63 | -0.06 | -0.01 | 0.12 | -0.15 |
| $P_M$ | -0.02 | -0.94 | -0.08 | 0.01 | 0.26 | -0.28 |
| $B_L$ | -0.13 | 0.16 | -0.07 | 0.20 | 0.10 | 0.10 |
| $B_M$ | -0.18 | -0.03 | 0.10 | 0.13 | 0.04 | 0.13 |
| $H_L$ | 0.07 | 0.28 | -0.05 | 0.11 | 0.12 | 0.18 |
| $H_M$ | 0.02 | 0.69 | -0.04 | 0.14 | 0.22 | 0.37 |

**Table S3**. Out-of-plane distortions in the PbRC protein environment of the dark structure for dataset b (Å).

| | saddling | ruffling | doming | waving | | propellering |
|---|---|---|---|---|---|---|
| | $B_{2u}$ | $B_{1u}$ | $A_{2u}$ | $E_{g(x)}$ | $E_{g(y)}$ | $A_{1u}$ |
| $P_L$ | 0.11 | -0.68 | -0.05 | -0.03 | 0.11 | -0.18 |
| $P_M$ | -0.08 | -0.91 | -0.10 | 0.01 | 0.22 | -0.28 |
| $B_L$ | -0.10 | 0.14 | -0.02 | 0.19 | 0.12 | 0.06 |
| $B_M$ | -0.09 | 0.05 | 0.04 | 0.14 | 0.10 | 0.09 |
| $H_L$ | 0.13 | 0.35 | -0.13 | 0.07 | 0.09 | 0.20 |
| $H_M$ | 0.08 | 0.57 | -0.11 | 0.16 | 0.20 | 0.32 |
